# Crystal structure of the first *de novo* designed phosphotriesterase ZAPP-1

**DOI:** 10.64898/2026.11.17.767279

**Authors:** Seth M. Woodbury, Donghyo Kim, Johnny Mendoza, Alex Kang, Hannah Nguyen, Asim K. Bera, David Baker

## Abstract

Designing a *de novo* phosphotriesterase requires scaffolding a dinuclear metal center bound by six catalytic residues, including a post-translationally carbamylated lysine, in a pocket that must also bind a small, flexible organophosphate and orient it for hydrolysis. We recently met this challenge using RFdiffusion3 to design the first *de novo* phosphotriesterases, and biochemically characterized our first hit, ZAPP-1 [1]. Here we report two crystal structures of ZAPP-1 at 1.42 and 2.28 Å resolution. The crystal structures are nearly identical to the design model scaffold (C*α* RMSD = 0.69, 0.71, and 0.99 Å for three copies across two independent crystals), and the higher-resolution crystal assembles the dinuclear Zn(II) site in the designed pocket with carbamylated K16. The catalytic residues are near the designed geometry but demonstrate flexibility across the structures, albeit mostly rotameric (C*β* RMSD ≤ 0.8 Å for both copies in form 1). AlphaFold3 predicts the designed catalytic conformation with substrate bound and the observed crystallographic deviations without it, suggesting an induced-fit mechanism upon substrate binding. Together with prior biochemical characterization, there is strong evidence that ZAPP-1 functions as designed, validating the high precision achievable with our RFdiffusion3 design pipeline, but the structures suggest that future design should better preorganize the active site. The ability to design experimentally accurate and increasingly sophisticated enzymes directly from the computer should enable a new future of programmable catalysis and chemistry.

## 1 Introduction

RFdiffusion3 is a generative deep learning method for designing biopolymers across diverse tasks and constraints, such as generating enzyme scaffolds around an active site catalyzing a reaction [1]. Building on our recent advances in atomic active site scaffolding and metallohydrolase design [2–4], we used RFdiffusion3 to design *de novo* phosphotriesterases, enzymes that hydrolyze neurotoxic organophosphates with broad applications in environmental decontamination and medical counter-measures [1]. While previous studies have reported phosphotriester hydrolysis by designed peptide assemblies [5, 6], a *de novo* metalloprotein [7], redesigned native enzymes [8–10], and proteins assembled from natural enzyme fragments [11], our phosphotriesterases are, to our knowledge, the first *de novo* enzymes designed explicitly for phosphotriester hydrolysis by generating new protein scaffolds around the catalytic residue functional groups of the *Pseudomonas diminuta* phosphotri-esterase [1], a well-studied enzyme famous for its broad and efficient hydrolysis of phosphotriesters and several organophosphorus pesticides and nerve agents [12–17]. Designing *de novo* phosphotriesterases was a significant challenge due to the complexity of their active site, which required scaffolding six catalytic residues, including a post-translationally carbamylated lysine, to bind two metal cofactors in a pocket that also binds and positions the flexible organophosphate, paraoxon, for hydrolysis.

In our first design campaign with RFdiffusion3, we screened 96 designs and identified 4 hits from the same scaffold that catalyzed paraoxon hydrolysis, the best of which was named Zinc-Activated Paraoxon Phosphotriesterase-1 (ZAPP-1). Biochemical characterization of ZAPP-1 showed that adding sodium bicarbonate, which interconverts to CO_2_ for driving lysine carbamylation, increased catalytic efficiency (*k*_cat_*/K*_M_) primarily by increasing *k*_cat_; alanine substitution of each designed catalytic residue abolished or nearly abolished activity, and ethylation trapping followed by mass spectrometry identified the trapped lysine carbamate [1, 18]. These findings strongly supported the design but left unresolved how closely ZAPP-1 adopted the intended scaffold and active-site geometry. Here we report two crystal structures of ZAPP-1 and compare them with the design model and AlphaFold3 (AF3) predictions [19] to examine the structural realization of this catalytic architecture and enzyme scaffold.

## 2 Results

### 2.1 Crystal structures of ZAPP-1 closely match the design model

To investigate the experimental structure of ZAPP-1, we solved two crystal structures of the tagless enzyme, yielding two forms: form 1 at 1.42 Å resolution with two chains per asymmetric unit, grown with *p*-nitrophenyl phosphate (pNPP; PDB 39CY), and form 2 at 2.28 Å resolution grown without added pNPP (PDB 39CX). Both forms reproduce the designed predominantly helical scaffold with near-atomic accuracy to the design model (C*α* RMSD = 0.69 Å and 0.71 Å for form 1 chains A and B; C*α* RMSD = 0.99 Å for form 2; RMSDs reported after outlier rejection) (Fig. 1A–C, Tables 1 and 2). The deviations from the design are mostly in a *β*-hairpin lid (75–85), which closes over the designed pocket and leads into the helix bearing E88, H89, E92 and H93; it is ordered and closely matches the design in form 1 but swings up to 6.0 Å away with its tip unresolved in form 2 (Fig. 1C). Additional deviation can be observed at the loop carrying H170 (165–171), which is ordered and design-like in form 2 but is mobile in form 1, where the two chains place it 6.6 Å apart (Fig. 1A). Natural enzymes exhibit similar heterogeneity, with the canonical bacterial phosphotriesterase populating open and closed conformations that respectively admit substrate and preorganize the site for paraoxon hydrolysis [20], and the *β*-hairpin flap over the dinuclear zinc center of a related metallohydrolase is mobile when the site is empty and damped when it is occupied [21]. We further analyze chain A of each form below, as its active site is the better resolved.

**Table 1.**
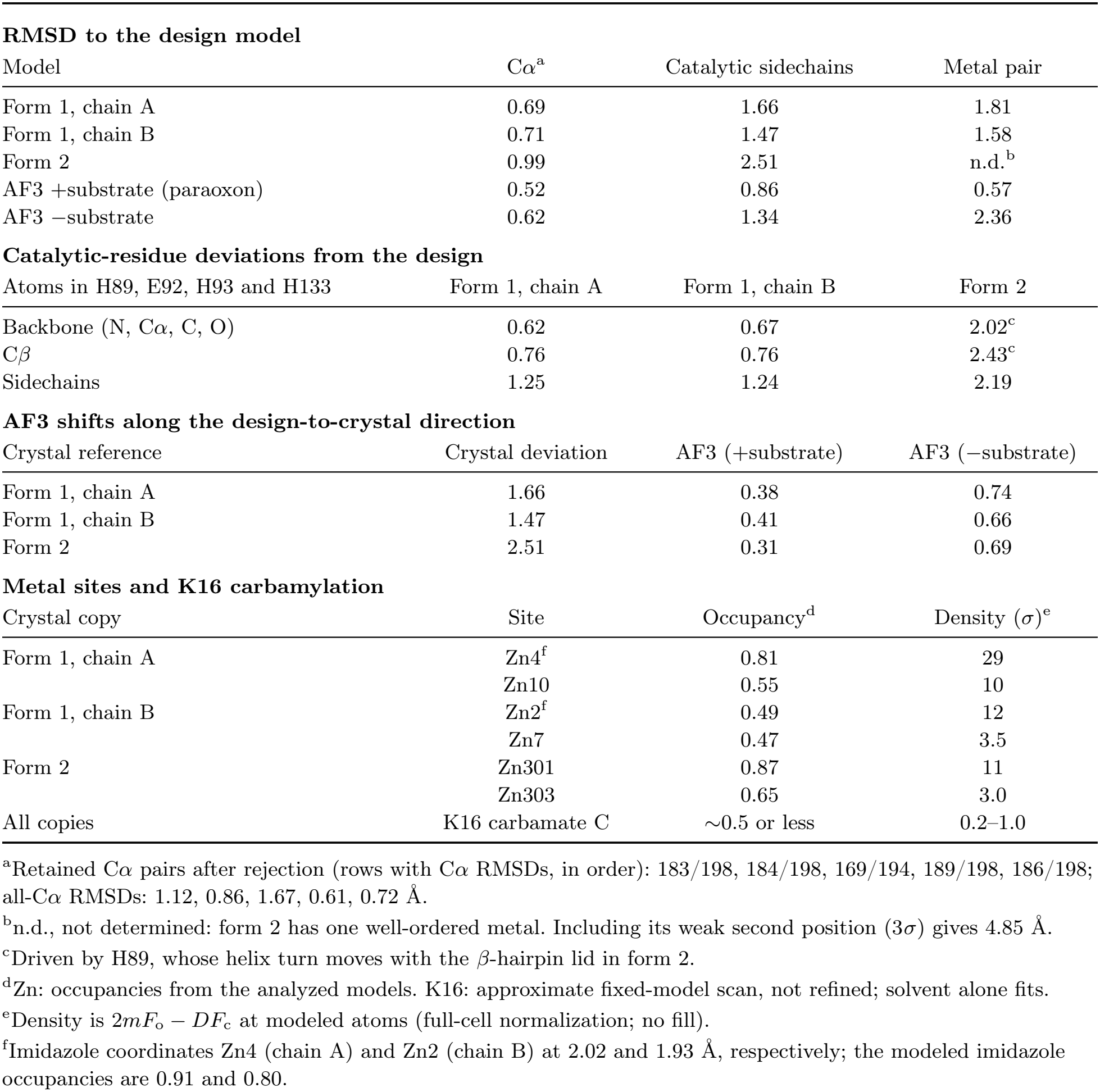
Structural comparisons of the ZAPP-1 crystal structures and AlphaFold3 predictions. RMSDs and shifts are in Å. Local values use the all-C*α* superposition without local refitting. Catalytic RMSDs and pooled directional shifts use 37 sidechain heavy atoms unless specified otherwise.

| **RMSD to the design model** |  |  |  |
| Model | Cα<sup>a</sup> | Catalytic sidechains | Metal pair |
|---|---|---|---|
| Form 1, chain A | 0.69 | 1.66 | 1.81 |
| Form 1, chain B | 0.71 | 1.47 | 1.58 |
| Form 2 | 0.99 | 2.51 | n.d.<sup>b</sup> |
| AF3 +substrate (paraoxon) | 0.52 | 0.86 | 0.57 |
| AF3 −substrate | 0.62 | 1.34 | 2.36 |
| **Catalytic-residue deviations from the design** |  |  |  |
| Atoms in H89, E92, H93 and H133 | Form 1, chain A | Form 1, chain B | Form 2 |
| Backbone (N, Cα, C, O) | 0.62 | 0.67 | 2.02<sup>c</sup> |
| Cβ | 0.76 | 0.76 | 2.43<sup>c</sup> |
| Sidechains | 1.25 | 1.24 | 2.19 |
| **AF3 shifts along the design-to-crystal direction** |  |  |  |
| Crystal reference | Crystal deviation | AF3 (+substrate) | AF3 (−substrate) |
| Form 1, chain A | 1.66 | 0.38 | 0.74 |
| Form 1, chain B | 1.47 | 0.41 | 0.66 |
| Form 2 | 2.51 | 0.31 | 0.69 |
| **Metal sites and K16 carbamylation** |  |  |  |
| Crystal copy | Site | Occupancy<sup>d</sup> | Density (σ)<sup>e</sup> |
| Form 1, chain A | Zn4<sup>f</sup> | 0.81 | 29 |
|  | Zn10 | 0.55 | 10 |
| Form 1, chain B | Zn2<sup>f</sup> | 0.49 | 12 |
|  | Zn7 | 0.47 | 3.5 |
| Form 2 | Zn301 | 0.87 | 11 |
|  | Zn303 | 0.65 | 3.0 |
| All copies | K16 carbamate C | ∼0.5 or less | 0.2–1.0 |

<sup>a</sup>Retained Cα pairs after rejection (rows with Cα RMSDs, in order): 183/198, 184/198, 169/194, 189/198, 186/198; all-Cα RMSDs: 1.12, 0.86, 1.67, 0.61, 0.72 Å.
<sup>b</sup>n.d., not determined: form 2 has one well-ordered metal. Including its weak second position (3σ) gives 4.85 Å.
<sup>c</sup>Driven by H89, whose helix turn moves with the β-hairpin lid in form 2.
<sup>d</sup>Zn: occupancies from the analyzed models. K16: approximate fixed-model scan, not refined; solvent alone fits.
<sup>e</sup>Density is 2*mF*<sub>o</sub> − *DF*<sub>c</sub> at modeled atoms (full-cell normalization; no fill).
<sup>f</sup>Imidazole coordinates Zn4 (chain A) and Zn2 (chain B) at 2.02 and 1.93 Å, respectively; the modeled imidazole occupancies are 0.91 and 0.80.

**Table 2.** Data collection and refinement statistics for the two ZAPP-1 crystal forms.

|  | Crystal form 1 | Crystal form 2 |
|---|---|---|
| PDB identifier | 39CY | 39CX |
|---|---|---|
| **Data collection** |  |  |
| Space group | *P* 1 2<sub>1</sub> 1 | *P* 2<sub>1</sub> 2<sub>1</sub> 2 |
| *a*, *b*, *c* (Å) | 68.419, 38.800, 72.396 | 63.836, 68.305, 37.175 |
| *α*, *β*, *γ* (°) | 90.000, 114.774, 90.000 | 90, 90, 90 |
| Resolution (Å) | 62.12–1.42 (1.45–1.42) | 68.30–2.28 (2.35–2.28) |
| *R*<sub>merge</sub> | 0.064 (1.267) | 0.258 (2.761) |
| *I*/*σ*(*I*) | 14.8 (1.5) | 6.9 (1.3) |
| Completeness (%) | 97.7 (95.5) | 99.8 (98.0) |
| Redundancy | 9.7 (7.0) | 13.1 (13.2) |
| CC<sub>1/2</sub> | 0.999 (0.562) | 0.996 (0.756) |
| **Refinement** |  |  |
| Resolution (Å) | 34.15–1.42 (1.45–1.42) | 46.64–2.28 (2.42–2.28) |
| Reflections | 63,225 (2,643) | 7,866 (1,263) |
| *R*<sub>work</sub> | 0.1826 (0.3041) | 0.2380 (0.3098) |
| *R*<sub>free</sub> | 0.2107 (0.3019) | 0.2811 (0.3413) |
| Protein atoms | 3,371 | 1,545 |
| Water atoms | 239 | 17 |
| Ligand atoms | 64 | 5 |
| Protein *B* (Å<sup>2</sup>) | 27 | 53 |
| Water *B* (Å<sup>2</sup>) | 34 | 47 |
| Ligand *B* (Å<sup>2</sup>) | 34 | 55 |
| Bond-length r.m.s. deviation (Å) | 0.017 | 0.018 |
| Bond-angle r.m.s. deviation (°) | 1.32 | 0.83 |

Values in parentheses refer to the highest-resolution shell; one crystal was used for each data collection.

**Figure 1.**
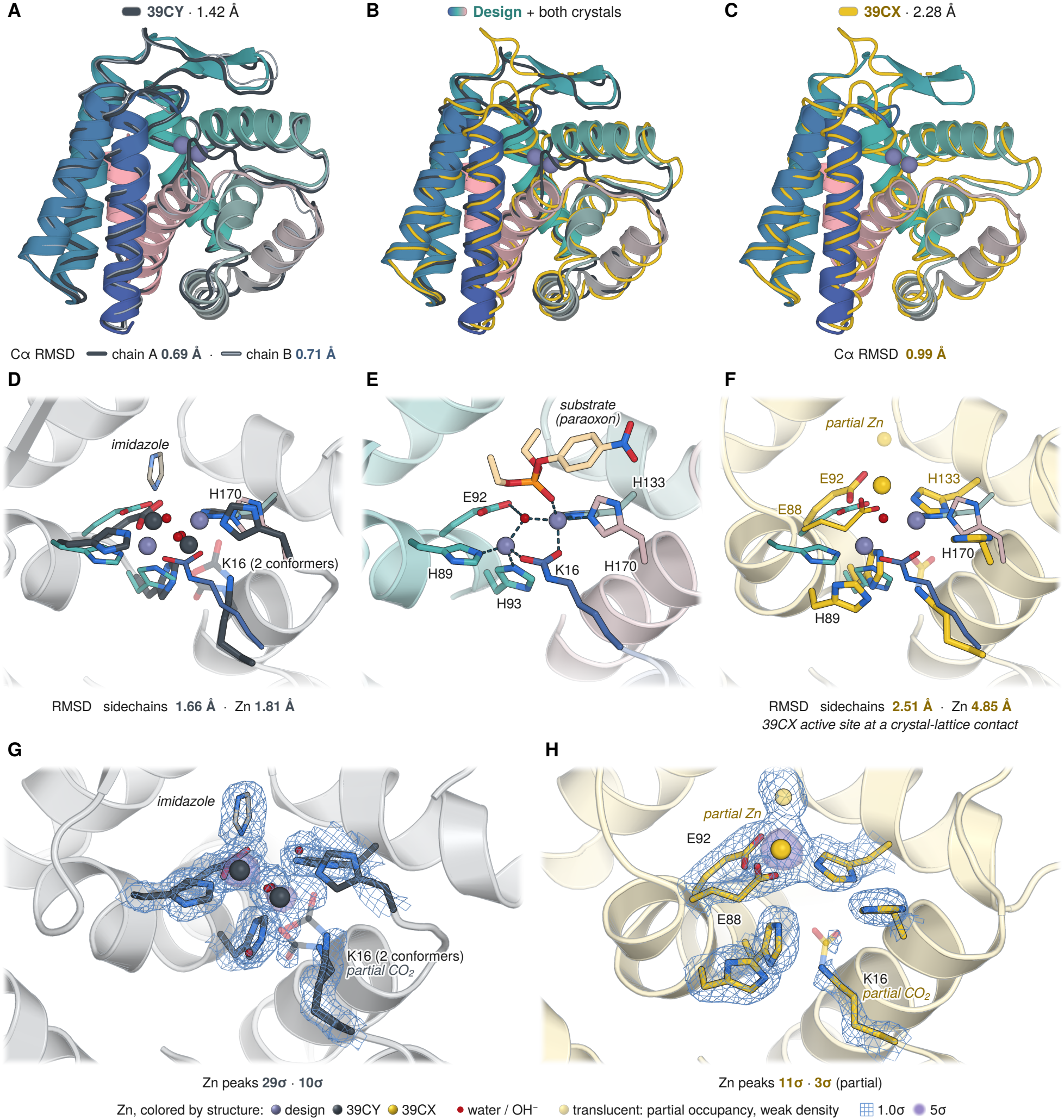
Crystal structures and RFdiffusion3 design model of ZAPP-1. (**A** to **C**) Crystal structures superposed on the RFdiffusion3 design model (ribbon colored blue to pink from the N to the C terminus); spheres mark the designed Zn(II) sites. (**A**) Form 1 (PDB 39CY, 1.42 Å), chain A as a graphite tube and chain B as a thin light trace. The copies align closely, with their H170-bearing loop (165–171) differing by up to 6.6 Å at corresponding C*α* positions in the design frame. (**B**) Design with chain A of both crystal forms. (**C**) Form 2 (PDB 39CX, 2.28 Å; saffron), with a displaced *β*-hairpin lid (75–85) and unresolved tip. Overlays use all shared C*α* atoms; the labeled C*α* RMSDs to the design use outlier-rejected fits (Table 1). (**D** to **F**) Active sites in a common view. In the overlays (**D** and **F**), the crystal backbones are pale gray or yellow ribbons; crystal sidechain carbons and Zn(II) spheres are graphite or saffron, respectively. In these overlays, the design is shown without its backbone, with catalytic sidechain carbons in the blue-to-pink gradient, Zn(II) spheres in lavender and the bridging hydroxide in red. (**D**) Form 1 chain A, showing both modeled K16 conformations and buffer imidazole coordinating Zn(II). (**E**) The designed catalytic site, with paraoxon, two Zn(II) ions bridged by hydroxide, and carbamylated K16 with H89, E92, H93, H133 and H170; dashes mark the designed metal coordination. (**F**) Form 2 chain A, with E88, E92 and H133 coordinating the well-ordered Zn(II). Sidechain RMSDs use the 37 catalytic sidechain heavy atoms, and metal RMSDs use the two modeled metal positions, after superposition on all shared C*α* atoms without local refitting. Form 2’s displayed 4.85 Å metal RMSD includes its weak second modeled site. (**G** and **H**) 2mFo–DFc electron density for form 1 chain A (**G**) and form 2 chain A (**H**), calculated without missing-reflection fill and contoured at 1.0*σ* (blue mesh) and 5.0*σ* around the metals (violet). Values below the panels are density heights at the modeled metal positions. Red spheres in the crystal views mark modeled water. Translucent atoms indicate weak density or modeled partial occupancy. K16 is modeled as carbamylated in both forms; a separate density analysis is consistent with partial carbamylation (Table 1), as expected for a reversible CO2 adduct, although solvent can also explain the density. The K16A knockout substitution has been shown to nearly abolish paraoxon-hydrolysis activity in ZAPP-1, and its lysine carbamate has been observed through mass spectrometry following ethylation trapping [1]. Form 2’s second metal site is modeled as partially occupied Zn(II) and has a density height of 3*σ*.

Neither form captures the complete catalytic geometry, but between them they recapitulate its separate parts (Fig. 1D–H). Form 1 assembles the dinuclear Zn(II) site in the designed pocket at 29*σ* and 10*σ*, with H89 and E92 coordinating the dominant metal as designed, and it recruits a buffer imidazole into the volume the substrate would occupy, showing that the site readily completes its coordination with an exogenous ligand (Fig. 1D, G). Form 2 holds a single well-ordered Zn(II) at 11*σ* through a different subset of catalytic residues, E92 and H133, and a designed residue, E88 (Fig. 1F, H). Form 1’s catalytic sidechains sit close to their designed positions, with backbone and C*β* RMSDs less than 0.8 Å from the design for the well-ordered catalytic residues in both copies and the remaining difference being mostly rotameric (Fig. 1D–F). K16 is modeled as carbamylated in both forms, in two conformations in form 1, and the density is consistent with partial carbamylation, as expected of a reversible CO_2_ adduct. Lysine carbamates are labile and routinely escape crystallographic detection: the catalytically essential carbamate of the class D *β*-lactamase OXA-10 was absent from its first structure [22] but was observed by ^13^C NMR in solution under conditions in which refined models lack it [23]. Since the modification was established in Rubisco on biochemical grounds alone [24], CO_2_ dependence of turnover rather than density has been the primary evidence for a functional carbamate, and ZAPP-1’s carbamate rests on its bicarbonate-dependent *k*_cat_ and on chemical trapping with the Meerwein reagent triethyloxonium for mass-spectrometry detection [1, 18]. The dinuclear Zn(II) site, the complementary portions of the designed coordination sphere recapitulated across the two forms, the exogenous ligand filling part of the substrate volume, and the density consistent with partial carbamylation are strong supporting evidence that the designed catalytic conformation is realized through induced-fit upon substrate binding.

### 2.2 AlphaFold3 supports induced-fit upon substrate binding

To further test the induced-fit hypothesis, we asked whether AF3 could recapitulate the deviations we observe experimentally when the substrate is absent (Fig. 2) [19]. Predicted with substrate bound, ZAPP-1 reproduces the design closely (C*α* RMSD = 0.52 Å; catalytic sidechain RMSD = 0.86 Å; Zn(II) RMSD = 0.57 Å) (Fig. 2A). Predicted without substrate and with water, the scaffold is unchanged (C*α* RMSD = 0.62 Å) but the catalytic residues relax 1.34 Å away from the designed geometry, most of the way to the 1.47 to 2.51 Å spanned by the three crystal copies (Fig. 2B, E). The substrate-free prediction therefore releases the designed catalytic conformation *in silico* while leaving the fold intact, with the catalytic residues shifted toward their positions in the crystal structures (Fig. 2C–F). Projected onto the design-to-crystal vector, the substrate-free prediction moves the sidechains roughly twice as far toward each crystal copy compared to the substrate-bound prediction (moved by 0.74 Å vs 0.38 Å toward form 1 chain A, respectively) (Fig. 2F). Together, the models suggest that substrate binding may induce small, local rearrangement in the active site to achieve the designed catalytic geometry, as observed in many native enzymes.

**Figure 2.**
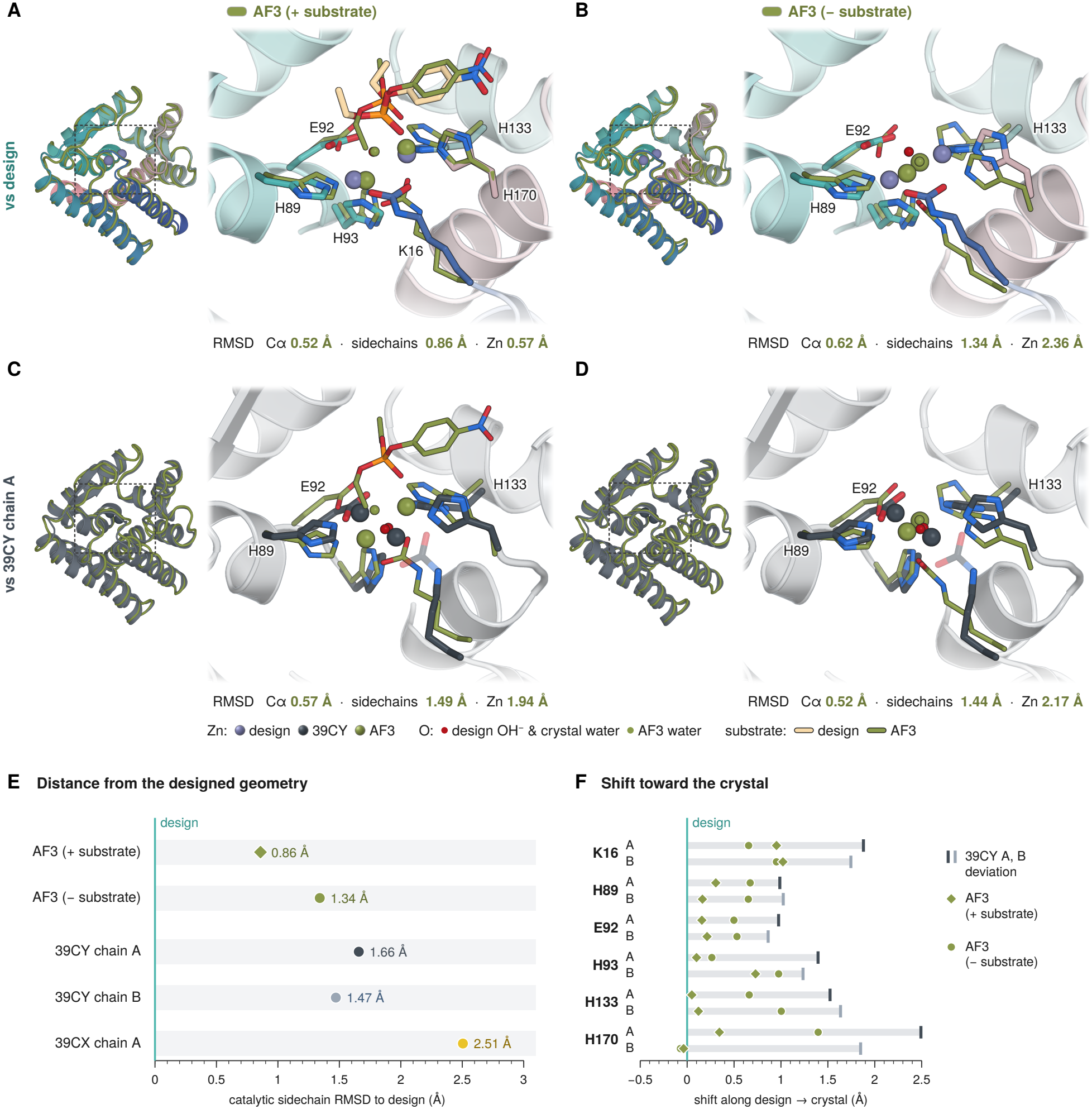
AlphaFold3 predictions of ZAPP-1 with and without substrate. (**A** to **D**) Representative AF3 predictions (olive) with two Zn(II) ions, a hydroxide ion, and the paraoxon substrate (“+ substrate”; **A, C**) or with two Zn(II) ions and water (“− substrate”; **B, D**), superposed on the design (blue to pink; **A, B**) or crystal form 1, chain A (graphite; **C, D**). Insets show the full-protein overlays, with boxes marking the enlarged active sites. Values below each panel are C*α* RMSD after outlier rejection, catalytic sidechain RMSD over 37 heavy atoms, and metalpair RMSD. The overlays and local RMSDs use all shared C*α* atoms without local refitting. Red spheres represent the catalytic water of the design model and the proposed catalytic water in the crystal; the AF3-predicted catalytic water is shown as an olive sphere. (**E**) Catalytic sidechain RMSDs to the design for the two AF3 predictions and three crystal copies. (**F**) For each catalytic residue, the AF3 sidechain displacement from the design projected along the corresponding design-to-crystal displacement, pooled over its heavy atoms after all-C*α* superposition. A and B identify the two form 1 chains. Gray bars end at each crystal copy’s sidechain RMSD to the design for that residue; diamonds mark the paraoxon-bound prediction and circles the water-containing substrate-free prediction. Positive values indicate a shift in the direction of the corresponding crystal structure.

## 3 Conclusion

The crystal structures of ZAPP-1 recapitulate the designed scaffold and assemble its dinuclear Zn(II) site with post-translationally carbamylated K16, hosting the catalytic residues at their designed backbone positions. Flexibility in the catalytic sidechains and their flanking structure resulted in each crystal form capturing different parts of the designed catalytic geometry, such that they collectively support it. Together with the alanine substitutions that abolish or nearly abolish activity at each catalytic residue, the bicarbonate dependence of catalysis, the lysine carbamate identified by chemical trapping [1, 18], and the AF3 predictions, the crystal structures support the design model but suggest that the designed catalytic geometry may be induced upon substrate binding. Future phosphotriesterase and enzyme design should buttress the catalytic residues more strongly, preorganizing the designed geometry rather than relying on substrate binding to form it [25, 26]. These structures provide an important confirmation of the accuracy of RFdiffusion3 for complex active site design in enzymes.

## Acknowledgments and funding

This work was supported by the National Science Foundation Graduate Research Fellowship Program grant DGE-2140004 (S.M.W.) and Defense Advanced Research Projects Agency grant HR00112520029, CFDA No. 12.910 (S.M.W., D.K., D.B.). Additional acknowledgments (S.M.W., D.B.) include the Defense Threat Reduction Agency; the Generative Unconstrained Intelligent Drug Engineering (GUIDE) program at Lawrence Livermore National Laboratory; and the U.S. Department of Energy’s Genesis Mission Phase II. We also acknowledge Schmidt Sciences, LLC (S.M.W., D.K., D.B.). Crystallographic work by A.K.B., A.K., H.N., and J.M. used resources 17-ID-1 and 17-ID-2 of the National Synchrotron Light Source II, a U.S. Department of Energy (DOE) Office of Science User Facility operated by Brookhaven National Laboratory under Contract No. DE-SC0012704. Beamtime was obtained through NECAT BAG proposal 317877. The Center for BioMolecular Structure is primarily supported by the National Institutes of Health, National Institute of General Medical Sciences, through grant P30GM133893, and by the DOE Office of Biological and Environmental Research under award KP1605010.

## Use of generative AI

Generative AI language tools were used in the preparation of this manuscript for LaTeX formatting, figure styling and assembly, coding assistance in the analysis, and copy-editing and review of drafts for grammar, internal consistency, and clarity. The study design, experimental work, data generation, analysis decisions, and scientific claims are the authors’ own. The authors are responsible for reviewing and verifying the full manuscript, including every reference, and take full responsibility for its contents.

## Author contributions

S.M.W. designed the phosphotriesterases. S.M.W. and D.K. experimentally characterized ZAPP-1 and prepared protein for crystallization screening. H.N. and A.K. screened crystallization conditions and prepared samples for diffraction data collection. J.M. and A.K.B. analyzed the diffraction data. D.B. and A.K.B. supervised the work. S.M.W. wrote the original draft and prepared the manuscript. All authors contributed to review and editing. S.M.W., D.K., J.M., A.K., H.N., and A.K.B. contributed equally.

## Data and materials availability

Coordinates and structure factors have been deposited in the Protein Data Bank under accessions 39CY (form 1, pNPP-supplemented crystallization) and 39CX (form 2, no added pNPP). The ZAPP-1 sequence is provided in Methods. Design and biochemical characterization are described in the published RFdiffusion3 manuscript (Butcher *et al*., “Atomic functional site scaffolding in biopolymers with RFdiffusion3,” *Science*, 2026) [1]. The RFdiffusion3 design and biochemical data are publicly available on GitHub at https://github.com/baker-laboratory/Metallohydrolase_Enzyme_Design. The accompanying crystallography package, comprising the ZAPP-1 design model, AF3 predictions and available inputs, crystal coordinate and reflection files used for the analyses, and structural analyses, is available at https://github.com/baker-laboratory/Metallohydrolase_Enzyme_Design/tree/main/Manuscript_Data/Phosphotriesterase_RFdiffusion3_Science_2026/eLetter_crystallography_addendum.

## 6 Methods

Materials and methods for cloning, protein expression, and purification were adapted from the published RFdiffusion3 manuscript [1]; see that report for additional details.

### 6.1 Materials

#### Solvents, buffers, and crystallization reagent

Milli-Q water was obtained in-house using a Q-POD ultrapure water dispenser (MilliporeSigma), and nuclease-free water was purchased from Invitrogen. HEPES, NaCl, ZnSO_4_, MgCl_2_, and NaHCO_3_ were purchased from MilliporeSigma. Buffers were filtered through 0.2-*µ*m PES membranes (Nalgene Rapid-Flow). *p*-Nitrophenyl phosphate disodium salt hexahydrate (pNPP; Calbiochem, product 4876) was obtained from MilliporeSigma.

#### Cloning and protein expression

The synthetic gene was ordered from IDT as an eBlock, and custom plasmid vectors were ordered from GenScript. BsaI-HFv2 (R3733L; 20 U/*µ*l), T4 DNA ligase (M0202L; 400 U/*µ*l), and 10× T4 DNA ligase reaction buffer were obtained from New England Biolabs, with the enzymes supplied separately; chemically competent BL21(DE3) cells were obtained from Intact Genomics. TB-II medium was purchased from MP Biomedicals, kanamycin sulfate from MilliporeSigma, and dioxane-free IPTG from Fisher Scientific. LB contained tryptone (Sigma-Aldrich), NaCl (MilliporeSigma), and Bacto Yeast Extract (Thermo Fisher Scientific). Glycerol was purchased from Sigma-Aldrich.

#### Protein purification

Lysozyme (L6876) and bovine-pancreas DNase I (DN255G) were purchased from Sigma-Aldrich, and Pierce protease inhibitor tablets from Thermo Fisher Scientific. Protein Select resin (17542102), the Superdex 75 Increase 10/300 GL column, and the ÄKTA pure system were from Cytiva. Amicon Ultra-15 centrifugal filters with a 10-kDa molecular-weight cutoff (MilliporeSigma, UFC901096) were supplied by Fisher Scientific.

### 6.2 Cloning and sequence verification

ZAPP-1 was produced using the tagless-protein workflow described for RFdiffusion3 [1]. An *Escherichia coli* -codon-optimized gene (IDT eBlock) was assembled into a custom T7 vector encoding an N-terminal His_6_–Protein Select fusion. For each 5 *µ*l Golden Gate assembly reaction, a master mix containing approximately 0.5 *µ*l vector (100 ng/*µ*l), 0.50 *µ*l 10× T4 DNA ligase reaction buffer (1× final; 50 mM Tris-HCl, pH 7.5, 10 mM MgCl_2_, 1 mM ATP, and 10 mM DTT), 0.15 *µ*l BsaI-HFv2 (20 U/*µ*l; 3 U), 0.10 *µ*l T4 DNA ligase (400 U/*µ*l; 40 U), and approximately 2.25 *µ*l RNase-free water was combined with approximately 1.5 *µ*l eBlock insert (10 ng/*µ*l). Water was adjusted as needed to bring each reaction to 5 *µ*l. Reactions were thermocycled for 30 cycles of 37 °C for 2 min and 16 °C for 2 min, followed by 60 °C for 10 min and a hold at 12 °C, with the heated lid set to 105 °C. One microliter of each assembly reaction was added to 6 *µ*l of chemically competent BL21(DE3) cells (Intact Genomics), and the cells were transformed by heat shock. Cells were recovered in SOC for 2 h at 37 °C and grown in LB with 50 *µ*g/ml kanamycin. Individual colonies were isolated, and 10 *µ*l of unpurified colony-PCR product amplified with T7 and T7-terminator primers was submitted to GENEWIZ (Azenta Life Sciences) for Sanger sequencing. Sequence-verified single-colony glycerol stocks were used for expression.

#### Expression construct

`MS–HHHHHH` (His_6_ tag)–SG–MVKIVSRKSLGVQNVYDIGVEKDHNFLLANGLIASN (Protein Select tag)`–SVG–`[design sequence]`–GS.`

**Design sequence (198 residues; numbering used for catalytic residues)**

~~~
1 ITEEQYRAIL EGLRE**K**QELI RKGEIPGAHP EVYELYEEIV RTLEEKGPTE EAIIEAVKVY LKKAKEIVEK 70
71 LADEEFEAPT GTKVTLAE**H**A L**EH**PAIALKA AGVELPPELK AALEKFDEIA RKLHPGLDAL RL**H**LAGIADD 140
141 PLFVELAREF GLGEDVERAR ASGFRISAA**H** GKGAFAVVFL FLYAIRKGYE DLIIEELK 198
~~~

**Expressed sequence (249 residues)**

~~~
1 MSHHHHHHSG MVKIVSRKSL GVQNVYDIGV EKDHNFLLAN GLIASNSVGI TEEQYRAILE GLRE**K**QELIR 70
71 KGEIPGAHPE VYELYEEIVR TLEEKGPTEE AIIEAVKVYL KKAKEIVEKL ADEEFEAPTG TKVTLAE**H**AL 140
141 **EH**PAIALKAA GVELPPELKA ALEKFDEIAR KLHPGLDALR L**H**LAGIADDP LFVELAREFG LGEDVERARA 210
211 SGFRISAA**H**G KGAFAVVFLF LYAIRKGYED LIIEELKGS 249
~~~

**Purified, intein-cleaved sequence (203 residues)**

~~~
1 SVGITEEQYR AILEGLRE**K**Q ELIRKGEIPG AHPEVYELYE EIVRTLEEKG PTEEAIIEAV KVYLKKAKEI 70
71 VEKLADEEFE APTGTKVTLA E**H**AL**EH**PAIA LKAAGVELPP ELKAALEKFD EIARKLHPGL DALRL**H**LAGI 140
141 ADDPLFVELA REFGLGEDVE RARASGFRIS AA**H**GKGAFAV VFLFLYAIRK GYEDLIIEEL KGS 203
~~~

Bold residues mark catalytic positions K16, H89, E92, H93, H133, and H170 (design numbering). Edge numbers refer to each displayed sequence. Design position 1 corresponds to product position 4, and K16 (product position 19) is encoded as lysine and modeled as KCX16 in the crystal structures.

### 6.3 Protein expression

Overnight 5-ml LB/kanamycin starters (37 °C, 225 rpm, 16–20 h) were used to inoculate 50-ml TB-II cultures (MP Biomedicals; 50 g/l) containing 50 *µ*g/ml kanamycin in 250-ml baffled flasks. A total of 600 ml culture was grown at 37 °C and 225 rpm to OD_600_ ≈ 0.6. Expression was induced with 1 mM IPTG and 200 *µ*M ZnSO_4_, followed by incubation at 18 °C and 225 rpm for 16–20 h. Cells were harvested at 4198 × *g* for 8 min at 25 °C and stored at −80 °C.

### 6.4 Tagless-protein purification

The base buffer was 40 mM HEPES, 50 mM NaCl, and 200 *µ*M ZnSO_4_, pH 8.0. Buffer components were from MilliporeSigma; buffers were prepared in Milli-Q water and filtered through 0.2-*µ*m PES membranes (Nalgene Rapid-Flow). Pellets were resuspended in 20 ml lysis buffer per 50 ml culture (240 ml total). Lysis buffer comprised base buffer with 1 mg/ml lysozyme and 10 *µ*g/ml DNase I (Sigma-Aldrich), 1 mM MgCl_2_, and one Pierce protease inhibitor tablet (Thermo Fisher Scientific) per 50 ml. Cells were sonicated on ice using a Qsonica Q500 with a four-tip horn (60% amplitude; 10 s on/20 s off; 180 s total on-time). Lysates were clarified at 14,000 × *g* for more than 45 min at 4 °C.

Clarified lysate was divided between two 12-ml Protein Select resin beds (Cytiva, 17542102; 120 ml lysate per bed), and the combined flow-through was applied to an additional approximately 20-ml bed. This split-intein system releases tagless protein without added protease or elution additives [27]. Each bed was washed with three 3.5-column-volume (CV) portions of base buffer. Column outlets were closed, approximately 3 ml buffer was left above each bed, and columns were incubated at room temperature for 20–24 h for on-column cleavage. Protein was eluted with three 1-CV portions of base buffer, yielding SVG–design sequence–GS. Eluates were concentrated to approximately 6.6 ml, clarified by centrifugation, and purified on a Superdex 75 Increase 10/300 GL column using an ÄKTA pure system (Cytiva) in base buffer.

Monomeric fractions were supplemented to approximately 45 mM NaHCO_3_ (MilliporeSigma) using a 500 mM stock prepared in Milli-Q water and incubated with shaking for approximately 2 h at room temperature. Protein was concentrated at 2000 × *g* to approximately 1.4 ml at 60 mg/ml using Amicon Ultra-15 centrifugal filters with a 10-kDa molecular-weight cutoff (MilliporeSigma, UFC901096; supplied by Fisher Scientific). Protein mass and tag removal were confirmed by intact-protein ESI-MS as described previously [1].

### 6.5 Sample preparation for crystallization

For co-crystallization of form 1 (39CY), a pNPP stock was prepared by dissolving 7.8 mg pNPP in approximately 520 *µ*l base buffer (approximately 40 mM). A 50-*µ*l portion of this stock was added dropwise to a 400-*µ*l aliquot of the purified protein solution with gentle mixing. The resulting 450 *µ*l sample contained approximately 53 mg/ml protein and 4.5 mM pNPP (approximately 2:1 pNPP:protein) and was clarified at 5000 × *g* for 1 min. For crystal form 2 (39CX), 1 ml of the 60 mg/ml protein stock was clarified at 4000 × *g* for 1 min and submitted without added pNPP. pNPP is a phosphomonoester and is distinct from paraoxon, the phosphotriester substrate used to characterize ZAPP-1 [1].

### 6.6 Crystallization and structure determination

All crystallization experiments were conducted using sitting-drop vapor diffusion. Crystallization plates were set up with 200-nl drops in 96-well format at 20 °C using a Mosquito LCP (SPT Labtech), then imaged using UVEX microscopes and a UVEX PS-256 (JAN Scientific). Diffraction-quality crystals of form 1 (39CY), co-crystallized with pNPP (53 mg/ml protein) at a 1:1 protein:reservoir ratio, formed in 0.07 M calcium acetate hydrate, 0.1 M imidazole, pH 7.0, and 28% (v/v) PEG 300, matching condition D3 of the LMB Crystallization Screen. For form 2 (39CX), the protein preparation without added pNPP (60 mg/ml) was screened at protein:reservoir ratios of 1:1, 1:2, and 2:1. Diffraction-quality crystals formed in 0.09 M Halogens Mix (sodium fluoride, sodium bromide, and sodium iodide), 0.1 M Buffer System 1 (imidazole and MES), pH 6.5, and 30% (v/v) Precipitant Mix 1 (40% PEG 500 MME and 20% PEG 20000), matching condition B1 of the Morpheus screen. All crystals were harvested directly from the crystallization drops and flash-frozen in liquid nitrogen.

Diffraction data were collected at beamlines 17-ID-1 (AMX) and 17-ID-2 (FMX) of the National Synchrotron Light Source II, using one crystal for each data collection. X-ray data for form 2 were evaluated and integrated using xia2/DIALS [28, 29], and data for form 1 using xia2.multiplex [30]; data were scaled and merged using Pointless/Aimless in the CCP4 program suite [31]. Starting phases for structure determination were obtained by molecular replacement using DIMPLE with the design model [32]. Following molecular replacement, iterative model building and corrections were performed manually in Coot [33], and refinement was carried out using REFMAC5 and CCP4 Cloud Refmacat [34–36]. Initial refinement was conducted using BUSTER to address Ramachandran, rotamer, and density-fit outliers, with refinement to convergence [37]. PDB-REDO was used to assess model quality between refinement rounds and to improve rotamers and density fit [38]. Final refinement was carried out in Phenix, with manual model building in Coot [39]. Model quality was evaluated using MolProbity [40]. Data collection and refinement statistics are recorded in Table 2. Coordinates and structure factors have been deposited in the Protein Data Bank under accession codes 39CY and 39CX.

### 6.7 Structural comparisons

Catalytic positions K16, H89, E92, H93, H133, and H170 were taken from the design model’s REMARK 666 records. Reported C*α* RMSDs were calculated with PyMOL align using polymer C*α* atoms and default outlier rejection (cutoff 2.0, up to five cycles). This procedure retained 183/198 atom pairs for form 1 chain A and 169/194 for form 2. A separate unweighted least-squares fit used every shared C*α* position of the design sequence without rejection; this fit defines the displayed superpositions and all local displacement measurements. Without rejection, the C*α* RMSDs are 1.12, 0.86, and 1.67 Å for form 1 chain A, form 1 chain B, and form 2, respectively (Table 1). Form 1 has coordinates for all 198 positions; form 2 lacks residues 78–81. No local refitting was applied to the active sites. Figures 1 and 2 use chain A for the detailed crystal comparisons; both copies of form 1 are included in the quantitative analyses. Each AF3 model in Fig. 2 was independently fitted to its indicated reference.

Catalytic sidechain RMSD includes 37 heavy atoms: eight in carbamylated Lys16, six in each histidine, and five in Glu92. Backbone atoms and hydrogens are excluded. The design’s separately represented carbamyl group was mapped to KCX atoms, and symmetry-equivalent carboxylate and carbamyl oxygens were assigned to minimize RMSD. Histidine nitrogen identities were retained.

The higher-occupancy Lys16 conformer was used for form 1 chain A. Omitting the three carbamate atoms gives sidechain RMSDs of 1.65, 1.49, and 2.55 Å for form 1 chain A, form 1 chain B, and form 2, respectively. The backbone, C*β*, and sidechain comparisons of four well-ordered catalytic residues in Table 1 use H89, E92, H93, and H133. Zinc-pair RMSDs were calculated after protein fit-ting, minimizing over the two possible metal correspondences without fitting the metals themselves. Additional zinc sites elsewhere in the asymmetric unit were excluded.

### 6.8 Local flexibility and crystal packing

Secondary structure was assigned with PyMOL. Per-residue deviations from the design were computed after superposition on all shared C*α* atoms and checked using a fit restricted to residues 30–70 and 86–198. The C*α* RMSD over residues 30–70 is 0.49, 0.46, and 0.75 Å for form 1 chain A, form 1 chain B, and form 2, respectively. Atomic *B* factors were expressed relative to each structure’s own mean, avoiding comparisons of absolute values between datasets at different resolutions. Packing contacts were identified with a symmetry-aware neighbor search using a 4 Å cutoff.

The *β*-hairpin lid in form 1 has *B* factors near its structure’s mean, and 13 of the 19 residues spanning positions 71–89 contact the other chain in the asymmetric unit. In form 2, residues 78– 81 are unmodeled, and the remaining lid has *B* factors approximately 1.5 times the mean, rising to 1.8 times the mean in the first strand. For residues 163–171, the corresponding ratios are 1.9 and 2.2 in the two copies of form 1 and 0.85 in form 2. The two modeled active-site metals are 2.87 Å apart in form 1 chain A and 2.30 Å apart in chain B, compared with approximately 3.3–3.6 Å in carbamylated-lysine-bridged dinuclear hydrolases [41–43]. These short separations and partial occupancies are consistent with alternative states rather than two fully occupied metal positions.

### 6.9 AlphaFold3 predictions and displacement analysis

The substrate-complex AF3 coordinates were supplied from the RFdiffusion3 figure workflow and contain a custom Zn/O component in addition to the other zinc and substrate. The original prediction input for this model was unavailable. The RFdiffusion3 holo-prediction protocol used Zn(II)-bound hydroxide [1]; the supplied Zn/O component is described accordingly, although its protonation is not specified in the coordinates. Substrate-free predictions used the 198-residue design sequence with KCX16, two zinc ions, and water. Input MSA and template fields were empty. Five supplied samples from seed 1 were analyzed. Sample 0 was selected for display as the lowest index among tied highest reported ranking scores. These samples represent within-run variation, not independent replicates. All five substrate-free samples were compared with the design and crystal structures using the same shared-position, untrimmed C*α* fit and 37-atom catalytic sidechain definition.

For a selected set of *N* sidechain heavy atoms, let **D, P**, and **C** be the concatenated 3*N* -component coordinates of the design, AF3 prediction, and crystal, respectively. After superposition on all shared C*α* atoms, the signed shift along the design-to-crystal direction was calculated as

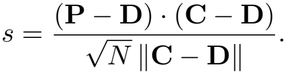

The normalization places the shift on the RMSD length scale; the crystal’s RMSD from the design is 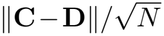. Positive values indicate a pooled displacement in the crystal direction. For Figure 2F, *N* is the number of heavy atoms in one catalytic sidechain and the calculation is performed for both form 1 chains. For Table 1, *N* = 37 and all three crystal copies are used. Both report the paraoxon-containing prediction and the displayed water sample. H89 and E92 are the best-defined catalytic residues. H170 shifts toward form 1 chain A but not chain B, whereas much of the H133 displacement is off-axis. Pooled over the 37 atoms, the substrate-bound and displayed substrate-free predictions have similar RMSDs to form 1 chain A (1.49 and 1.44 Å, respectively), motivating the residue-resolved and directional comparison. The five substrate-free water predictions have sidechain RMSDs of 1.23–1.34 Å to the design.

### 6.10 Electron-density maps and carbamate occupancy

Electron-density maps were calculated from the available 2*mF*_o_ − *DF*_c_ coefficients without missing-reflection fill and normalized over each full unit cell. Quoted map heights are values at the modeled atom positions. Maps were transformed together with the corresponding crystal model. The figure density views use 1.0*σ* blue meshes masked within 1.8 Å of the displayed atoms, and translucent violet 5.0*σ* surfaces masked within 1.6 Å of the modeled zinc positions. Both modeled Lys16 conformations are shown for form 1 chain A. Molecular views were ray-traced in PyMOL and assembled with Arial labels. Site and density views were composited from separate rendering passes for the molecular representations, without changing atom coordinates or map values.

Carbamate occupancy was estimated by a scan without re-refinement. Structure factors for the crystal models used in the analysis were recomputed with GEMMI [36], with bulk-solvent correction and anisotropic scaling. With the scale and solvent mask fixed, the model was decomposed into the structure without the carbamate, a variable carbamate-carbon contribution, and a variable contribution from the two carbamate oxygens. Carbon and oxygen occupancies were scanned relative to the modeled lysine occupancy, with CE and NZ fixed. The scan was evaluated using *mF*_o_ − *DF*_c_ density at the atoms, the mean squared difference density within 1.6 Å of the carbamate atoms, and local real-space correlation and residuals. Two displacement-parameter treatments were compared: the values in the analyzed models and values matched to NZ. The E92 carboxylate was scanned as a calibration control. Carbon-based estimates provide the carbamate occupancy summary in Table 1; these are estimates from fixed-model scans, not newly refined occupancies. The scan permits partial carbamate plus solvent but does not distinguish it uniquely from solvent at the oxygen positions.

